# Making Sense of Neural Networks in the Light of Evolutionary Optimization

**DOI:** 10.1101/2023.11.27.568922

**Authors:** Anton V. Sinitskiy

## Abstract

To what extent can evolution be considered as the sole first principle that explains all properties of nervous systems? This paper proposes an innovative, mathematically rigorous perspective on understanding nervous systems from an evolutionary perspective, leveraging methods of nonequilibrium statistical physics. This approach allows for modeling an exhaustive evolutionary optimization of nervous systems irrespective of any specific molecular and cellular mechanisms, approximate neuronal models or evolutionary history events. This novel method may shed light on key properties of biological neural networks and may also have potential relevance for understanding artificial neural networks.

## Introduction

“Nothing in biology makes sense,” Theodosius Dobzhansky wrote in 1973, “except in the light of evolution.”^1^ In this work, we focus on a particular case of that nothing: a nervous system. Can we say that evolution explains all properties of nervous systems, or, in other words, that all properties of nervous systems can be strictly derived from evolution as the sole first principle?

The literature has extensively discussed nervous systems as an evolutionary adaptation, and many properties of nervous systems (degree of complexity, plasticity, modularity, tuning of sensory systems, etc.) have been explained within this framework.^2-11^ However, such investigations are often limited to verbal reasoning; in those cases when derivations are mathematically formalized, they rarely do exhaustive search in the space of all hypothetically possible variants, being restricted to a comparison of a limited number of alternatives. For example, an optimal shape of a neuronal spike was studied by varying numerical parameters in a particular (e.g., Hodgkin–Huxley) model of a neuron,^12,13^ but, to the best of our knowledge, have never been investigated in the most general form, namely: What is the optimal shape of a neuronal spike in general, regardless of any approximate models that we use to describe neurons, or molecular mechanisms that known-to-us neurons rely upon for functioning? In this work, we attempt to set the stage for an exhaustive search for evolutionarily optimal solutions for nervous systems, not limited by any approximate models of neurons, neuronal networks, sensors, effectors, etc., or specific molecular and cellular mechanisms that underlie the functioning of the nervous system.

Until recently, a general mathematical solution to this problem had been impossible. The situation has changed with the development of a new approach, based on the methods of nonequilibrium statistical physics. This approach links dynamic equations of a system – in this case, the nervous system – to two entities. The first one is the nonequilibrium steady state probability density or, equivalently, a potential function (also called a potential energy function), which is in one-to-one correspondence to this density. The second entity is the so-called solenoidal or rotation field (or its equivalents, see Methods). So far, this approach has been used to analyze the dynamics of fixed neural networks.^14-26^ For the first time, to the best of our knowledge, we recognize that this approach is more powerful than previously thought. It can be used for an exhaustive evolutionary optimization of the properties of neurons and neural networks.^27,28^

We hypothesize that the key properties of nervous systems and biological neural networks can be understood and predicted by solving the mathematical problem of the optimization of the evolutionary fitness as a functional of the two above-mentioned functions. In this paper, we outline what a mathematical apparatus of such a theory might look like and discuss possible complications along this path.

Theoretical results on biological neural networks presented in this work may also be relevant for artificial neural networks.^29^ Due to the generality of the system’s description, insensitive to specific molecular mechanisms of neurobiological phenomena, it may be extended in a large degree to non-biological systems, using the language of nonequilibrium statistical physics and thermodynamics. Artificial neural networks, like biological ones, are dissipative systems. Their working cost, perhaps, could be defined based on general physical concepts, such as the entropy production rate. On the other hand, there must be a significant difference between these two classes of networks. Biological neural networks are optimal from the viewpoint of evolutionary fitness, while artificial networks are intelligently designed based on other goal functions, including their economic efficiency. A comparative analysis of these two types of networks might clarify the similarity and dissimilarity of their properties and make extrapolations from biological evolution to the development of artificial intelligence and, possibly, artificial general intelligence more justified.

## Methods

### Nonequilibrium statistical physics approach to neuronal dynamics

In this subsection, we introduce the existing approach, highlighting important aspects that are rarely emphasized in prior literature. Consider a stochastic model of the nervous system described by *N* dynamic variables, *x*_*j*_(*t*), which depend on time *t*. In the simplest case, each neuron is described by a single variable *x*_*j*_, interpreted as its membrane potential. Then, the rate of change of the potential of each neuron depends on the current values of its own potential, the potentials of other neurons and the external (sensory) inputs to neurons, which explicitly depend on time:

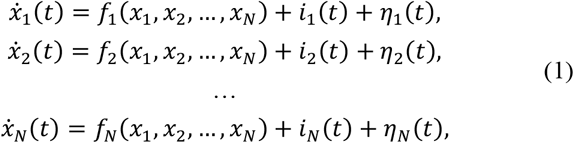

where *f*_*j*_ describe the internal dynamics of the nervous system, *i*_*j*_(*t*) represent the external input (in units of potential), and *η*_*j*_(*t*) are random variables (noise). Note that equations (1) can also be applied to the case when each neuron is described by several variables *x*_*j*_, for example, as in the Hodgkin–Huxley model.^30^

We complement this model with a model of the surrounding world, further termed “the environment”:

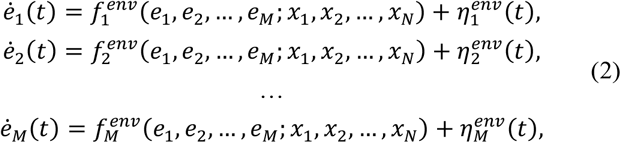

where *e*_*j*_ are variables describing the state of the external world, *M* is the total number of such variables, 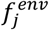 are functions describing the change in the external world depending on its current state (*e*_1_, …, *e*_*M*_), as well as on the current state of the modeled nervoussystem (*x*_1_, …, *x*_*N*_) to account for the nervous system’s active influence on the external world via effectors, and 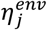 are random variables (noise). Note that dynamic equations for the environment (or the impact of the nervous system on it) that include second-order and higher-order derivatives can be written in the general form of equation (2) using a trick well-known from classical mechanics, namely by increasing the number of independent variables (e.g., introducing momenta as independent variables in addition to coordinates).

Further, sensory input into the nervous system depends on the state of the environment (and, possibly, the current state of the nervous system):

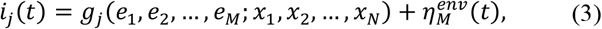

where *g*_*j*_ are functions showing how the state of the external environment (and, possibly, of the nervous system) determines sensory input, 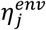 is noise in sensory input, and index *j* runs from 1 to *N*. In a vector form, equations (1), (2) and (3) can be written as:

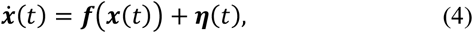

where the vector ***x*** encodes the state of the whole system (the nervous system and the environment) at a given moment in time *t*:

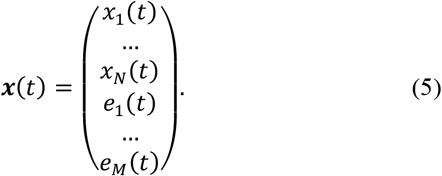

the vector function ***f***(***x***) is composed of the functions 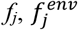 and *g*_*j*_ introduced above, and similarly ***η***(*t*) is a vector build from 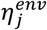and 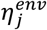. Unlike original equations (1), equation (4) describes the dynamics of the system in a closed form, and the function ***f***, unlike *i*_*j*_(*t*) above, does not explicitly depend on time. From this perspective, the nervous system is the result of a spontaneous symmetry breaking into two sets of components of ***x*** (namely, variables *x*_*j*_ and *e*_*j*_) in a dynamic universe. Only considering the full system, characterized by a set of variables describing the nervous system and the external world (including changes in it caused by the nervous system), provides a full picture of the neural activity. Unlike previous work,^16,19,26^ such a full description allows us to avoid the need to artificially set external input for each neuron and its explicit time dependence [*i*_*j*_(*t*)], and it allows us to avoid assumptions about *i*_*j*_(*t*), such as their constancy over time or nullification. Also, a complete description that includes the external world with its laws avoids one-sided consideration of neural networks only as devices for perceiving signals from the outside world. In previous work, neural networks only explained the world in different ways, but the point is to change it.

Regarding the random variables ***η***(*t*), the standard assumptions^14,18,19,21,22,26,31,32^ seem reasonable that they follow Gaussian distributions with zero mean values (otherwise non-zero means could be included in the definition of ***f***) and covariance

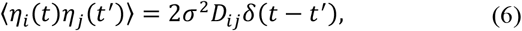

where *σ*^2^ sets the scale of random fluctuations common for all variables ***x*** (in similar stochastic equations in physics, *σ*^2^ is interpreted as the temperature in units of energy), *δ*(·) is the Dirac delta function, and *D*_*ij*_ in this work are assumed independent of ***x*** and diagonal (*D*_*i,j≠i*_ = 0). These two conditions on *D*_*ij*_ serve only to simplify the derivations and in principle can be lifted. Note that *σ*^2^ could be absorbed into the definition of *D*_*ij*_, and it is not done so only to have a single small parameter for perturbative expansion later in the analysis.

Now we move from considering the individual trajectory of one instance of the system to considering the probability distribution of trajectories of the ensemble of systems. Let *P*(***x***,*t*) be the probability density of such distribution, that is, the probability of finding the given system at time *t* with values of the variables in the range from ***x*** to ***x+dx*** equals *P*(***x***,*t*)*dx*_1_*…dx*_*N*_*de*_1_*…de*_*M*_. Then the change in *P* over time is described by the Fokker-Planck equation:

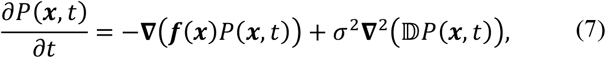

where ∇ is the gradient vector

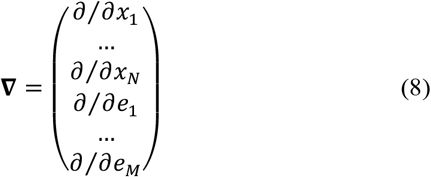

and the diffusion matrix ⅅ is formed by the elements *D*_*ij*_. This equation can be rewritten in terms of the flux ***J*** defined as

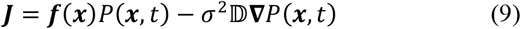

in the following way:

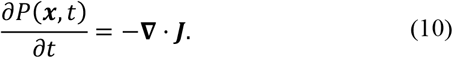

If the probability distribution over time reaches a stationary (unchanging) state, then the probability density in such a state *P*_*stat*_(***x***) is determined by the condition:

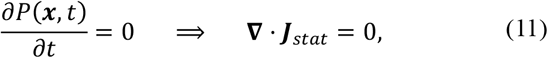

where

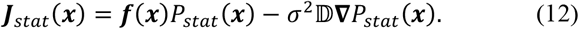

As shown in the Appendix, equation (11) implies that ***J***_*stat*_ can be written in general as

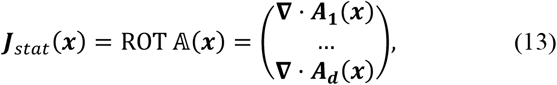

where ***A***_*i*_(***x***) are rows of an anti-symmetric matrix 𝔸(***x***) called a rotation potential, and ROT is a multidimensional generalization of the common three-dimensional curl operator.^33^ Therefore, the dynamics of the whole system [namely, the functions forming the vector ***f***(***x***)] can be parameterized, as follows from equations (12) and (13), in terms of *P*_*stat*_(***x***) and 𝔸(***x***):

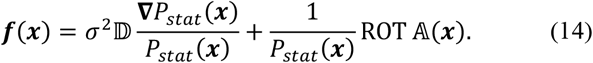

It is convenient to introduce a potential energy function *u*(***x***) by analogy with the Boltzmann distribution in statistical physics:

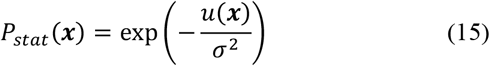

[recall that *σ*^2^ is an analog of the temperature in units of energy; the partition function is absorbed here into an additive constant in the definition of *u*(***x***)]. Then equation (14) transforms to

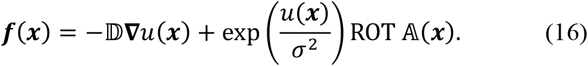

Consider its expansion in terms of *σ*^2^ as a small parameter. The left-hand side of this equation is independent of *σ*^2^. To make the right-hand side well-behaved at small *σ*^2^, introduce a new anti-symmetric matrix function ℚ(***x***) defined by

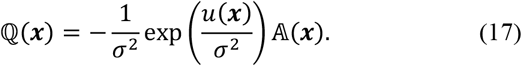

Then, equation (16) transforms into

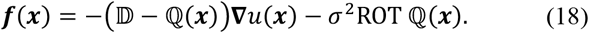

Expanding *u*(***x***) and ℚ(***x***) in Taylor series in terms of *σ*^2^:

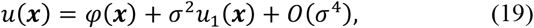

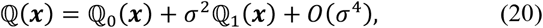

and plugging these expansions into the right-hand side of equation (18), we get a Taylor series in terms of *σ*^2^. Equating the terms with the same order in *σ*^2^, we get a sequence of equations, starting from the following equation for the zeroth-order terms:^15,18,20,21,23-25,32,34,35^

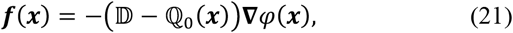

continuing to the next term, of the order of *σ*^2^, that writes as

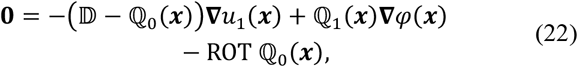

etc. Note that both equations (18) and (21) are exact and provide a decomposition of ***f***(***x***) into a gradient and solenoidal (rotational) components, but in the existing literature only equation (21) is discussed (under the names of the Helmholtz, Helmholtz-Ao or Helmholtz–Hodge decomposition^15,17,18,20,21,24,35^). This is presumably because this equation seems simpler, due to the absence of the ROT term with derivatives of the elements of ℚ in it. However, this simplification comes at a price of using the leading (in *σ*^2^) terms *φ*(***x***) and ℚ_0_(***x***) instead of the exact functions *u*(***x***) and ℚ(***x***). If computing the stationary state probability density *P*_*stat*_(***x***) alongside ***f***(***x***) is required, as in our subsequent analysis, then instead of dealing with the ROT term, one will have to build the Taylor series for *u*(***x***), as depicted in equation (19). This means solving iterative equations like equation (22) to determine *u*_1_(***x***), ℚ_1_(***x***) and higher-order terms, a task that can be challenging.

We argue that using equation (21) and *φ*(***x***) instead of *u*(***x***) may be problematic. Adopting such a replacement presumes a limit of *σ*^2^ → 0. Yet, in this limit, the stationary state probability density converges to a Dirac delta function, because:

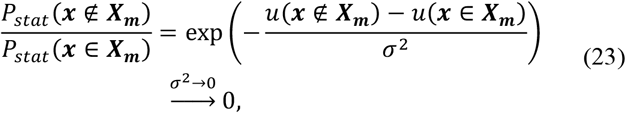

where ***X***_***m***_ is the subset of ***x*** values at which *P*_*stat*_(***x***) reaches its minimal value,

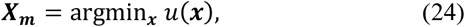

since by construction

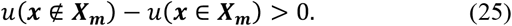

Furthermore, to ensure that *P*_*stat*_(***x***) remains normalized, *P*_*stat*_(***x*** ∈ ***X***_***m***_) must approach infinity as this limit is reached. Thus, *P*_*stat*_(***x***) reduces to a Dirac delta function in the limit of *σ*^2^ → 0. Hence, strictly speaking, the weak noise assumption provides information only about the most probable stationary state of the system and its immediate vicinity. It does not apply even to local minima of the potential energy, contradicting the practice of applications of this approach in neural network analyses.^14-26^ This general conclusion is supported by specific artifacts of the weak noise assumption revealed in a simple model of a nervous system that we reported earlier.^27,28^

### Formulation of the problem of evolutionary optimization

We now apply the above approach, previously used to describe nonequilibrium systems in constant neural networks,^14-26^ to a new problem, namely, evolutionary optimization of neural networks. The nervous system, though requiring organisms to expend additional resources, ensures greater population survivability. The evolutionarily optimal properties of nervous systems are determined by the balance between the evolutionary benefit of the nervous system and its cost. Let us formalize expressions for these factors.

#### Probability of an organism’s death

The death rate of organisms (the probability of death per unit of time) *P*_*death*_, assuming ergodicity, can be expressed as follows:

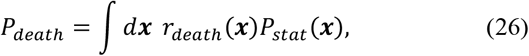

where *r*_*death*_(***x***) is the probability density of death per unit time given that the whole system (both the environment and the nervous system) are in the state ***x***, and the integration is performed over all possible values of ***x***. For example, the probability of being eaten by a predator may depend on the distance between the organism and the predators, that is, on some of the environmental variables, which, without the loss of generality, we choose as (*e*_1_, …, *e*_*P*_), where *P* ≤ *M*. Then, if the probability of being eaten is the same for all close distances, then

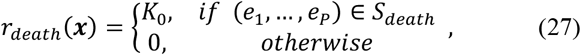

where *K*_0_ is a constant having the units of inverse time, and *S*_*death*_ is the subset of the values of distances (*e*_1_, …, *e*_*P*_) at which the organism is eaten. Then the expression for the death rate simplifies to

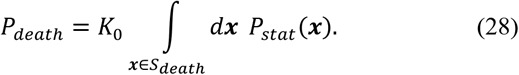

This estimate is based on the assumption that the stationary distribution exists and extends to the region *S*_*death*_, where death occurs with a certain probability, though not with certainty. An alternative approach may involve extending the Fokker-Planck equation for the case of guaranteed death of organisms at the border of *S*_*death*_, which may require an explicit solution of a complicated differential equation. We find this alternative approach unnecessarily complex.

#### Cost of the work of the nervous system

In generating an action potential, a neuron loses some of its ions (for example, K^+^), while additional amounts of other ions (for example, Na^+^ or Cl^−^) penetrate the cell. To maintain the neuron’s ability to generate action potentials in the future, ions need to be pumped against their electrochemical potentials.^5,7,13,36-38^ As we argued elsewhere,^28^ the average cost of the ion pumps’ work in a given neuron per unit time is proportional to the average value over time of |*x*_*j*_(*t*) ™ *x*_*j*,0_|, where *x*_*j*_ is one of the components of the vector ***x***, namely the membrane potential of the given neuron, and *x*_*j*,0_ is the resting potential of this neuron. Assuming ergodicity of the system, the average over time can be replaced by the average over the ensemble, therefore, the cost of the working nervous system is on average proportional to the value of *I*_1_, defined as

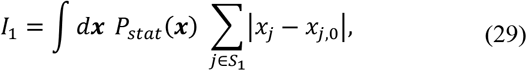

where *S*_1_ is the set of all indices for which *x*_*j*_ is the membrane potential of a neuron in the nervous system [as stated above, a neuron may be described by some of the variables *x*_*j*_ from equations (1), while other variables *x*_*j*_ may refer to other physical values or hidden variables, not the membrane potentials]. This expression for the cost includes the working cost of sensors. The mechanism of sensor action is such that ion channels sensitive to an external signal (presence of certain chemicals, light, mechanical pressure, temperature, etc.) open, and the membrane potential of the sensor neuron changes due to the passage of ions through the ion channels. As with the generation of a spike in an interneuron, the expenditure of free energy is associated with pumping ions against gradients of their electrochemical potentials, and therefore, it is included in the calculation above, as long as the membrane potentials of sensory cells are included in the set *S*_1_.

#### Cost of the work of effectors

Equation (29), however, does not include the working cost of effectors (for example, cilia or muscles that allow the organism to move in space). In a particular case of a simple model we studied previously^27,28^ (with dynamic variables that we denote here as *x*_1_ and *e*_1_), the cost was expressed in terms of the difference Δ*f*_2_ between *f*_2_, which is one of the components of ***f***(***x***), interpreted as the speed at which the distance from an organism to the nearest predator changes, and the movement speed of the predator itself *f*_*pred*_(*e*_1_), which is an externally set function that is not affected by the evolutionary optimization of the nervous system of the given organism:

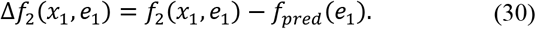

The cost of the working effectors in that example was proportional to the average of 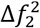over time, which we replaced, assuming ergodicity, by the ensemble average. As a result, this component of the cost was found to be proportional to *I*_2_, defined as^28^

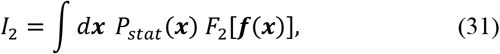

with 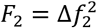 in that particular case. The quadratic dependence appears from the fact that the work performed during mechanical movement equals the product of the force *F* and the speed *v*. When moving in a viscous medium, the movement speed is proportional to the force, *v* ∼ *F*, which means that the work per unit time is proportional to the square of the speed *v*^2^. We do not have a general expression for *I*_2_ applicable to all neural networks yet, but we expect that equation (31) be valid in many, if not all, cases, given the freedom to choose specific functional forms for *F*_2_. Moreover, in many cases *F*_2_ may assume the following form:

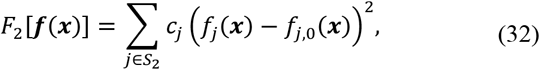

where *S*_2_ is the set of all indices for which *x*_*j*_ is the variable associated with effectors changing distances in the system, *c*_*j*_ are positive constants, *f*_*j*_ is the *j*-th component of ***f***(***x***), and *f*_*j*,0_ are some reference functions determined by the physical properties of the whole system or only the environment. A general expression for *I*_2_ remains unknown yet. One possible strategy to find it would be to generalize specific equations referring to various species and their effectors. Another solution might be based on general considerations about thermodynamics of dissipative systems, as discussed below.

#### Possible use of general nonequilibrium thermodynamic equations to compute the cost of the neural network

Equations (29) and (31) have a rather general form, but they were derived with a reference to certain material mechanisms in the nervous system and its effectors. Nonequilibrium thermodynamics offers a different approach to this problem, providing general mechanism-insensitive expressions that may turn out to be relevant for a general theory of neural networks.^16,18,19,23,31,32,39-43^ A nervous system is dissipative and, as any dissipative system, its thermodynamic cost can be expressed as the rate of entropy production 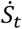, which can be calculated in the steady state as

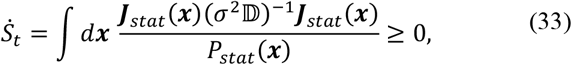

where the steady state flux ***J***_*stat*_ was defined by equation (12). Taking into account equation (18), the steady state flux can be rewritten as

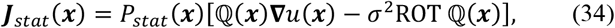

and the rate of entropy production as

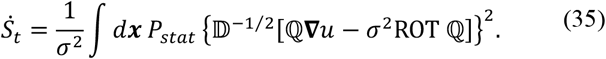

As expected, in the absence of the solenoidal potential (ℚ = **0**) the rate of entropy production equals zero.

The question of whether this expression for 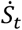 can lead to equations (29) and (31) as particular cases needs further investigation.

#### Evolutionary optimality in general

The death of some of the organisms reduces the growth rate of the population by the amount of *P*_*death*_ given by equation (26). The work of the nervous system and effectors, while reducing *P*_*death*_, also requires the organisms to spend their resources, which can no longer be directed towards population growth, thus reducing the rate of the population growth by amounts proportional to *I*_1_ and *I*_2_. Thus, the population growth rate *R* can be written as:

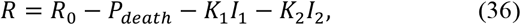

where *R*_0_ represents the contributions to the population growth rate determined by other factors not related to the nervous system (for example, the availability of food) and that we consider constant within the framework of this model, and *K*_1_ and *K*_2_ are positive constants translating the cost of the nervous system from the units of voltage and velocity squared for *I*_1_ and *I*_2_, respectively, first into the units of energy per unit time and then to the units of the population growth rate.

Natural selection optimizes ***f***(***x***) among functions that are physically and biologically achievable, thereby providing the maximum value of *R*, or equivalently, minimizing the functional *I* defined as

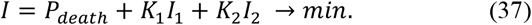

The values of *P*_*death*_, *I*_1_ and *I*_2_ are all non-negative (typically, positive), therefore:

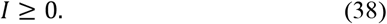

A direct minimization of *I* as a functional of ***f*** is technically complicated and can hardly be done without extensive numerical simulations for each specific system of interest. This explains why modeling an exhaustive optimization of neural networks has not been previously performed. This difficulty can now be circumvented due to the representation of ***f*** in terms of *u*(***x***) and ℚ(***x***) by equation (18), and *P*_*stat*_ in terms of *u*(***x***) by equation (15). With these recent results, the functional *I* can be considered as a relatively simple functional of *u* and ℚ:

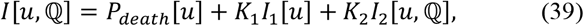

which may allow for an analytical solution in a general case:

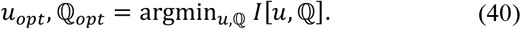

Then, the optimal solution for the dynamic laws governing the neural network *regardless of its material implementation or historical accidents* can be found from the optimal value of ***f***:

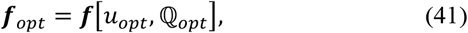

as defined by equation (18).

In general, when minimizing ***I***[***u***, ℚ], we must impose certain restrictions on the permissible values of *u* and ℚ, reflecting physically and biologically possible functions ***f***. In particular, the structure of the external physical world cannot be changed by natural selection acting on the nervous system of the organism of interest (we neglect indirect effects caused by large-scale ecological processes). Therefore, the components of ***f*** that refer to the environmental degrees of freedom [*e*_1_, …, *e*_*M*_ in equations (2) and (5)] with the nervous system not working [*x*_*j*_(*t*) = *x*_*j*,0_, that is, the resting potentials for variables in the set *S*_1_ as defined above, and some other reference constant values for non-voltage variables] should be restricted by external physical factors:

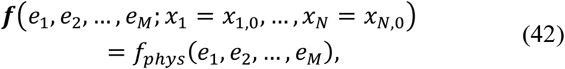

where *f*_*phys*_ are some functions reflecting the physical laws governing the environment. Also, the probability distribution function *P*_*stat*_ should be normalized to unity, implying another restriction on *u*:

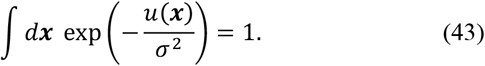

Other restrictions may be revealed as we better understand this approach and apply it to various specific cases.

## Results

The current formulation of the variational problem, as dictated by equations (39)-(41) and constrained by (42) and (43), remains incomplete. This was revealed when analyzing a simplified two-dimensional model, which adheres to the aforementioned general constraints.^27,28^ It turned out that a representation for 𝔸 (or, equivalently, ℚ) can be constructed such that condition (42) holds for all values of ***x***, not just those linked to a resting nervous system. Consequently, the *I*_2_ term in the functional *I*, which is the only term that depends on ℚ as per equation (39), always equals zero. However, this solution is neither biologically nor physically plausible as it suggests the absence of a motor response in the organism. Thus, to address this inconsistency, there is a need for more comprehensive constraints on the minimization of ***I***[***u***, ℚ] (see Discussion below for more detail).

Moreover, within this basic model, we derived a spiking nature of the nervous system as a consequence of evolutionary optimization.^27,28^ To sidestep the challenges with the foundational principles, we chose the potential *u* in a form that corresponds to ***f*** with converging stationary distributions and reduced the minimization of *I* as a functional to minimization of *I* as a function of several numerical parameters. Immediately, the spike-like behavior over the neuronal degree of freedom followed from a minimization of *K*_1_*I*_1_+*K*_2_*I*_2_, the part of *I* that depended on this degree of freedom.^27,28^ This derivation of a spike-like behavior of neurons should be generalizable to the general case considered in this work. It is worth noting that when considering equations (29) and (31) [in the latter case, taking into account equations (32) and (42)] both integrals can become negligible if *P*_*stat*_ draws close to a product of Dirac delta functions over the neuronal degrees of freedom *x*_*j*_ included in the sets *S*_1_ or *S*_2_ as defined above. On the other hand, the other term in *I*, namely *P*_*death*_, is largely or entirely governed by environmental variables, ensuring that it does not infringe upon the partial minimization of *I* over the neuronal degrees of freedom included in the sets *S*_1_ or *S*_2_. Hence, the analogy with the simplest model reported previously^27,28^ should extend to the conclusion that *P*_*stat*_ factorizes into terms including delta functions over the degrees of freedom in the sets *S*_1_ or *S*_2_, therefore, the potential *u* along these dimensions in ***x*** should be infinitely (in the limit) steep, and the dynamical equations should include terms returning the corresponding dynamic variables *x*_*j*_ to their resting values infinitely (in the limit) fast, that is, demonstrating a spike-like dynamic behavior along these neuronal degrees of freedom.

Lastly, our study provides a generalization of the Helmholtz decomposition^15,17,18,20,21,24,35^ to a finite *σ*^2^ parameter. As demonstrated above, equations (23)-(25), relying solely on a weak noise limit (*σ*^2^ → 0), as commonly done,^15,18,20,21,23-25,32,34,35^ can sometimes produce misleading outcomes even at local minima of the potential energy, let alone the regular functioning of neural networks. In addition, we have previously provided specific examples of artifacts that stem from neglecting finite *σ*^2^ effects in the toy model, namely the divergence of a component of ***f*** at the biologically most plausible values of its arguments, and a loss of an arbitrary contribution to *Q* (the only independent component of ℚ in that toy model) not restricted by the choice of the other component of ***f***.^27,28^ Taken together, these results on the weak noise limit highlight the need for a more rigorous mathematical approach in the transition from the Fokker-Planck representation to the representation of a system in terms of the functions *u* and ℚ.

## Discussion

Our study introduces a novel approach for understanding neuronal networks from a theoretical perspective. By leveraging the evolutionary principle and distancing ourselves from specific molecular and cellular mechanisms or the evolutionary history, we aim to offer insights into the intricacies of neural networks.

The cornerstone of our method blends the established nonequilibrium thermodynamics formalism, which expresses the temporal dynamics of systems using time-invariant entities like the potential *u* and the rotation potential 𝔸 (or ℚ uniquely defined by 𝔸), with evolutionary optimization. In this narrative, *u* and ℚ are no longer static entities but are independent functions that capture the possible dynamics of the system and therefore account, among other things, for mutations and variability. By changing the perspective from the ***f*** representation (describing the nervous system in terms of specific ion channels with a given kinetics, etc.) to the representation in terms of *u* and 𝔸, we get enabled to write explicit equations for the functional *I* to be optimized by evolution, and perform this variational optimization explicitly.

Several other distinctions also set our approach apart from prior research. The functional to be optimized is biologically motivated and directly linked to the population growth rate, unlike previously proposed loss functions like the sum of squared differences of outputs (membrane potentials of neurons).^44-48^ Unlike previous studies,^18,21,22,32^ we do not assume ℚ to be constant. The set of dynamic variables includes not only the neuronal degrees of freedom, but also those of the environment, lifting the restriction of stationary input to a neural network.^16,19,26^

Recently, an extension of the free energy principle to natural selection has been proposed.^35^ However, it is not clear to us how this extension could technically be applied, for example, to our simple model discussed above.^27, 28^ Moreover, this extension relies on the Helmholtz–Hodge decomposition, which, as we demonstrated earlier, implies a questionable weak noise limit, and on the Markov blankets assumption, which, to the best of our knowledge, has never been derived, either precisely or approximately, from fundamental physical principles.

Our approach, though currently in its initial phase, can potentially shed light on certain characteristic properties of neural networks, irrespective of their material manifestation. This encompasses the sharp excitation and relaxation responses of neurons, the energetic efficiency of neural system operations after evolutionary optimization, and the connection between global and local optimization strategies (higher population growth rate vs. faster kinetics and greater sensitivity of ion channels), as outlined for the general case in this work, and demonstrated previously for a particular simple case.^27,28^

However, challenges remain. The framework, as it currently stands, is not fully comprehensive. We have identified gaps in our understanding, especially when it comes to general restrictions on *u* and ℚ, which hinder a purely *ab initio* solution to the evolutionary optimization problem. Also, generalized expressions for the working cost of the nervous system are not clear. In particular, a general form of *F*_2_ from equation (31) is not known yet. Further generalization to multidimensional cases may reveal new issues.

Potential avenues for further work, which are not mutually exclusive, may include the following. The problem of searching for additional principles to be included into the evolutionary optimization of *I* needs further investigation. Necessary and sufficient constraints on *u* and ℚ might be introduced by demanding that a stationary solution to the Fokker-Planck equation *P*_*stat*_ exists for dynamic equations (4) with ***f*** defined from *u* and ℚ by equation (18). The analysis of the toy model^27,28^ demonstrates that the mere existence of *u* with necessary properties like condition (43) is not sufficient for the existence of *P*_*stat*_, despite equation (15). With some values of ℚ (for example, the above-mentioned unacceptable solution for the toy model) the stationary distribution may diverge.

Additional constraints on *u* and ℚ might formalize physical and biological limitations on the permissible functions ***f*** in real neurons. In particular, it seems intuitively plausible that expressions for ***f*** may separate contributions from variables (*x*_1_,…, *x*_*N*_) standing for internal neural dynamics and (*e*_1_, …, *e*_*M*_) responsible for sensory input, which, however, is not automatically ensured in the general case given by equations (4), (18) and (41). Due to the presence of terms involving ℚ in equation (18), separation of variables in ***f*** does not imply their separation in *u*(***x***), and vice versa. It remains to be clarified whether such a separation of variables in ***f*** is a consequence of physical limitations or a property of an optimal solution, invariant to the material implementation of the nervous system.

Additional terms might be added to the functional (39) optimized by evolution. Our estimates of all three quantities *P*_*death*_, *I*_1_ and *I*_2_ are based on certain approximations. It is possible that we did not include some qualitatively different components of the cost of the nervous system operation, which, in particular, do not allow achieving Δ*f*_2_ = 0 as a general solution, as well as additional terms in the accounted components of the cost, which become prominent during the minimization of *I*_1_ and *I*_2_ and their approach to zero.

An intriguing question is the relationship between the components of the cost of the nervous system given by equations (29) and (31) and the general expression for the “cost” of a non-equilibrium dissipative system (33). Can the terms *I*_1_ and *I*_2_ be derived as special cases of the general expression for the entropy production rate? Or does entropy production represent another component of the nervous system’s cost? (Note that entropy production is quadratic in ℚ, which could shift the optimal value of ℚ and thereby solve the problem of zeroing Δ*f*_2_). The question of the relationship between 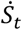 and ***K***_1_***I***_1_ + ***K***_2_***I***_2_ is not trivial. In particular, in the simple model mentioned above, in the limit of infinite distance between the organism and the predator, the minimization of 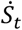 should not impose any constraints on the amplitude of neuronal excitation, because the problem splits into two separate one-dimensional problems, and a one-dimensional system is not dissipative, and thus should not lead to restrictions about the sharp character of neuron excitation, and instead allows arbitrary values of the membrane potential in this long-distance limit.

Finally, we note that the proposed theoretical framework may be extended to apply not only to biological, but also artificial neural networks. If it is possible to generalize the expressions for the cost of biological neural networks, as given by equations (29) and (31), to an arbitrary nonequilibrium system – perhaps, but not necessarily, through the entropy production rate as given by equation (33) – then a variational cost minimization problem, similar to the variational problem given by equation (37), may be formulated. This approach might yield new insights into both the nervous system and artificial intelligence systems, including potential limitations on emerging artificial general intelligence, which has recently been attracting much public attention.

In this study, we presented a novel theoretical framework to analyze neuronal networks, anchored in evolutionary principles and the nonequilibrium thermodynamics formalism. Moving away from conventional models, our approach provides a fresh perspective, unhindered by the specificities of molecular and cellular mechanisms or by random events in evolutionary trajectories. While we recognize our model’s limitations, it offers significant advancements, highlighting the need for further refinement. As with other theories, our proposal may undergo refinements, akin to the evolution of Bohr’s contradictory atomic model. The most promising future directions may involve exploring the relationship between neural system costs and entropy production rate and the potential extension of our approach to understanding artificial neural networks.

## Appendix

A generalization of the Helmholtz decomposition in a space with an arbitrary number of dimensions (not necessarily 3) leads to the statement that a *d*-dimensional (with *d = M + N* to agree with the main text) vector field ***J***(***x***) can be represented as a sum of two terms:^33^

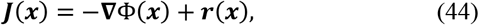

where the first term ™∇Φ(***x***) is an irrotational gradient field, which can be written as the negative gradient of the gradient potential Φ(***x***) (unlike the cited paper, we use the physical convention, with the negative sign before the gradient), and ***r***(***x***) is a solenoidal (divergence-free) rotation field. A sufficient (but not necessary) condition for equation (44) to hold is that ***J*** ∈ *C*^2^(ℝ^*d*^, ℝ^*d*^) and decays faster than |***x***|^-*δ*^ for |***x***| → ∞ with *δ* > 2. It seems plausible that these conditions are satisfied, taken into account equation (9), because the probability distribution of trajectories should rapidly decay for too strong neuronal excitations or a physical dissociation of the system into its parts.

Equation (44) can be inverted to express Φ(***x***) and ***r***(***x***) in terms of ***J***(***x***). In particular, the scalar potential Φ(***x***) can be found as

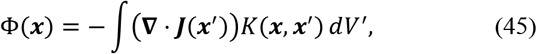

where the integration is performed over the *d*-dimensional space of all possible values of variables ***x’***, the integration kernel *K*(***x***,***x’***) is the fundamental solution of Laplace’s equation in *d*-dimensional space:

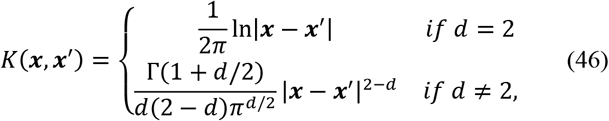

and Γ(·) is the gamma function.

The rotational field ***r***(***x***) in equation (44) can be written in terms of a rotation operator ROT, which is a *d*-dimensional generalization of the common three-dimensional curl operator, and a rotation potential 𝔸(***x***), which is a matrix-valued map (simply speaking, a matrix with elements depending on 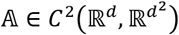 :

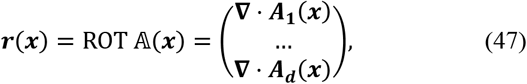

where ***A***_*i*_(***x***) are *d*-dimensional vectors formed by elements *A*_*ij*_(***x***) of 𝔸(***x***) with *j* = 1, …, *d*, and 𝔸(***x***) is anti-symmetric:

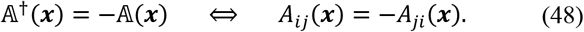

For the stationary state flux ***J***_*stat*_, taking into account equations (11) and (45), we conclude that the gradient field term in the generalized Helmholtz decomposition vanishes, and only the rotational field term, given by equation (47), is present, leading to equation (13) in the main text.

